# Extrasynaptic localization is essential for α5GABA_A_ receptor modulation of dopamine system function

**DOI:** 10.1101/2023.07.12.548744

**Authors:** Alexandra M. McCoy, Thomas D. Prevot, Md Yeunus Mian, Dishary Sharmin, Adeeba N. Ahmad, James M. Cook, Etienne L. Sibille, Daniel J. Lodge

## Abstract

Dopamine system dysfunction, observed in animal models with psychosis-like symptomatology, can be restored by targeting Gamma-Aminobutyric Acid type A receptors (GABA_A_R) containing the α5, but not α1, subunit in the ventral hippocampus (vHipp). The reason for this discrepancy in efficacy remains elusive; however, one key difference is that α1GABA_A_Rs are primarily located in the synapse, whereas α5GABA_A_Rs are mostly extrasynaptic. To test whether receptor location is responsible for this difference in efficacy, we injected a small interfering ribonucleic acid (siRNA) into the vHipp to knock down radixin, a scaffolding protein that holds α5GABA_A_Rs in the extrasynaptic space. We then administered GL-II-73, a positive allosteric modulator of α5GABA_A_Rs (α5-PAM) known to reverse shock-induced deficits in dopamine system function, to determine if shifting α5GABA_A_Rs from the extrasynaptic space to the synapse would prevent the effects of α5-PAM on dopamine system function. As expected, knockdown of radixin significantly decreased radixin-associated α5GABA_A_Rs and increased the proportion of synaptic α5GABA_A_Rs, without changing the overall expression of α5GABA_A_Rs. Importantly, GL-II-73 was no longer able to modulate dopamine neuron activity in radixin-knockdown rats, indicating that the extrasynaptic localization of α5GABA_A_Rs is critical for hippocampal modulation of the dopamine system. These results may have important implications for clinical use of GL-II-73, as periods of high hippocampal activity appear to favor synaptic α5GABA_A_Rs, thus efficacy may be diminished in conditions where aberrant hippocampal activity is present.

**Significance Statement:** Dopamine activity is known to be altered in both psychosis patients and in animal models, with promising new antipsychotics restoring normal dopamine system function. One such drug is GL-II-73, a positive allosteric modulator of α5GABA_A_Rs (α5-PAM). Interestingly, previous research has shown that a positive allosteric modulator of α1GABA_A_Rs (α1-PAM) does not share this ability, even when directly given to the ventral hippocampus, a region known to modulate dopamine activity. One potential explanation for this difference we examined in this study is that α1GABA_A_Rs are primarily located in the synapse, whereas α5GABA_A_Rs are mostly extrasynaptic. Determining the mechanism of this differential efficacy could lead to the refinement of antipsychotic treatment and improve patient outcomes overall.

## 1. Introduction

Gamma-Aminobutyric Acid type A receptors containing the α5 subunit (α5GABA_A_Rs) have received considerable attention as a therapeutic target for multiple disorders involving hippocampal pathology, likely due to their enhanced expression within CA1 and CA3 regions of the hippocampus ^1–3^. Of particular interest to the pharmaceutical industry are positive allosteric modulators (PAMs) selective for α5GABA_A_Rs because of their low propensity for side effects compared to nonselective benzodiazepines, which are known to cause sedation through actions mediated by α1 subunits ^4,5^. Preclinical studies using α5-PAMs have demonstrated a range of beneficial effects when given acutely including: anxiolytic, antidepressant, and pro-cognitive effects ^6,7^. When administered chronically, α5-PAMs can reverse stress- or age-related neuronal atrophy in the hippocampus and prefrontal cortex ^7,8^. Additionally, α5-PAMs have shown promise in preclinical studies as antipsychotics ^9–11^. These results suggest that α5-PAMs may have therapeutic utility for multiple disorders, especially those in which aberrant hippocampal activity is present.

Though the dopamine hypothesis asserts that psychosis is driven by excessive dopamine, convergent evidence suggests that dopamine dysregulation is secondary to aberrant hippocampal output, which drives the dopamine system dysfunction ^12–14^. Indeed, increased hippocampal activity has been observed in humans with psychosis ^15^ and in rodent models ^12–14^. Further, we have previously demonstrated that decreasing hippocampal activity using pharmacological ^16^, cell-based ^17,18^, or surgical ^19^ approaches effectively normalizes dopamine system function and related behaviors in rodent models with schizophrenia-like symptomatology. Thus, we posit that augmenting the function of α5GABA_A_Rs in the ventral hippocampus (vHipp) will likely have the same effect and normalize dopamine system function and behavior in a stress-based model displaying psychosis-like pathology. Indeed, we have previously demonstrated that this is the case in animal models used to study both schizophrenia and post-traumatic stress disorder (PTSD) where aberrant dopamine system function is present ^10,11,20^. This evidence suggests that α5GABA_A_Rs may represent a viable therapeutic target for the treatment of psychosis across multiple disorders.

Interestingly, nonspecific positive allosteric modulation of GABA_A_Rs or selectively targeting hippocampal α1GABA_A_Rs are largely ineffective at reversing aberrant dopamine system function ^11,20^. One crucial difference between α5- and α1GABA_A_Rs is that α5GABA_A_Rs can dynamically travel between the extrasynaptic space, where they regulate tonic inhibition ^21–23^ and the synapse, where they mediate phasic inhibition ^24^, whereas α1GABA_A_Rs are almost exclusively synaptic ^25,26^. Unlike typical extrasynaptic receptors that are diffuse in the membrane, α5GABA_A_Rs form clusters ^25,27,28^.

This clustering is mediated through an interaction with radixin, a scaffolding protein that anchors the receptor to actin, concentrating receptors in the extrasynaptic space (**Figure 1;** Loebrich et al., 2006). The radixin-α5 interaction is phosphorylation-dependent, such that dephosphorylation of radixin decouples the two proteins, allowing the receptor to diffuse freely through the membrane ^27^. In mutant mice that express a phosphorylation incompetent radixin, α5GABA_A_Rs co-localize with gephyrin, the inhibitory synaptic scaffolding protein, suggesting that, in the absence of a radixin interaction, α5GABA_A_Rs will move into the synapse and interact with gephyrin ^27,28^. Furthermore, the shifting of α5GABA_A_Rs into the synapse appears to be physiologically relevant, as induction of long-term potentiation in the hippocampus can also increase synaptic relocalization of α5GABA_A_Rs ^24^. Indeed, it has been hypothesized that the purpose of α5GABA_A_Rs clustering is to serve as a readily releasable pool of GABA_A_Rs to rapidly adjust to perturbations of excitatory/inhibitory balance, with periods of high activity increasing the contribution of α5GABA_A_Rs to inhibitory post synaptic potentials ^24^.

**Figure 1.**
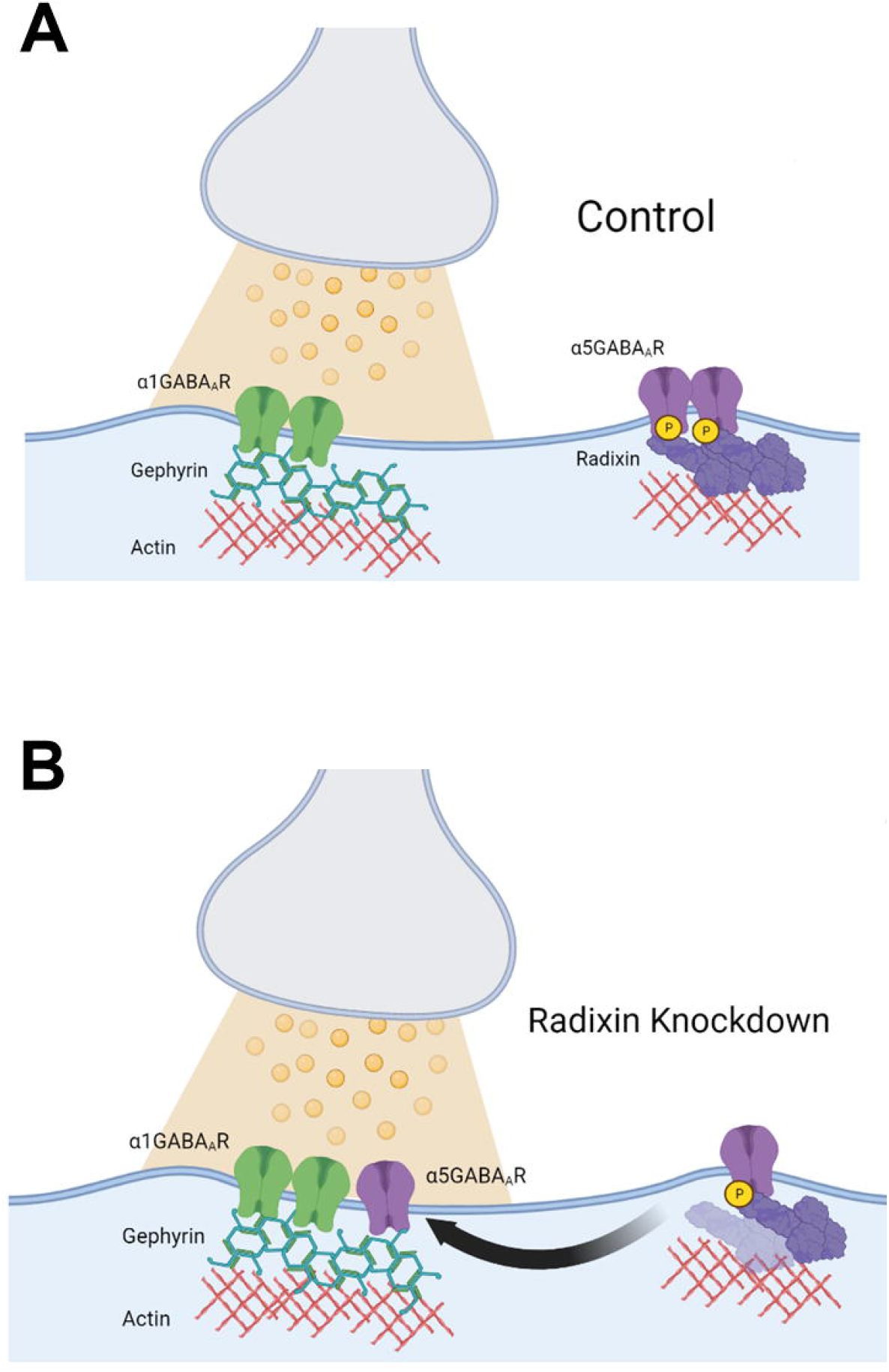
Schematic of radixin knockdown. Diagram of an inhibitory synapse and surrounding extrasynaptic area under (***A***) baseline conditions and (***B***) when radixin is knocked down. Figure made using Biorender.

Given the remarkable difference in antipsychotic-like efficacy between targeting α5- and α1-GABA_A_Rs ^11,20^, we sought to examine if receptor location (extrasynaptic vs synaptic) of α5GABA_A_Rs could influence the effects of a selective α5-PAM, GL-II-73, on dopamine system function and sensorimotor gating (prepulse inhibition of startle; PPI), a dopamine-dependent behavior often affected in psychosis ^29^. We injected small interfering RNA (siRNA) targeting radixin or a scrambled siRNA, as a control, directly into the vHipp of adult rats. Under these conditions, we examined the effects of GL-II-73 on dopamine neuron population activity in the ventral tegmental area (VTA) and on PPI. Exposure to an inescapable shock (IS) for two days is a validated model used to study PTSD-like pathology in rodents ^30^, a condition often comorbid with psychosis ^31^. In this model, we have demonstrated that the rats exhibit psychosis-like symptomatology, such as robust alterations in dopamine neuron activity and deficits in PPI ^32^ that can be reversed by GL-II-73 ^10^. Here, we now report that this reversal was blunted following radixin knockdown, causing α5GABA_A_Rs to shift into the synapse. These findings establish a clear relationship between α5GABA_A_R localization and the antipsychotic-like efficacy of GL-II-73. Such information is critical for clinical use of α5-PAMs, as α5GABA_A_R location appears to be activity dependent ^24^, and conditions in which hippocampal hyperactivity is present (e.g. epilepsy-induced psychosis) may promote a synaptic shift of α5GABA_A_Rs and would decrease antipsychotic efficacy in these individuals.

## 2. Materials and Methods

All experiments were performed in accordance with the guidelines outlined in the USPH Guide for the Care and Use of Laboratory Animals and were approved by the Institutional Animal Care and the Use Committees of UT Health San Antonio and U.S. Department of Veterans Affairs.

### 2.1. Animals

Adult, male (350-400g) and female (250-300g) Sprague Dawley rats purchased from Envigo (Indianapolis, IN, USA) were used for all experiments. Rats were kept on a 12 h light/dark cycle. Food and water were provided *ad libitum*. GL-II-73 or vehicle (85% H_2_0, 14% propylene glycol, 1% Tween80) were administered directly into the vHipp (100ng/uL; 0.75μL; AP⍰-5.3 mm, ML⍰±⍰5.0 mm, from Bregma DV −6.0⍰mm from brain surface) at a rate of approximately 0.5 μL/min 20 minutes prior to electrophysiology or behavior. This dose and timing was selected based on previous characterization ^6^ as well as our own data ^10,11^.

### 2.2 siRNA mediated knockdown of Radixin

Rats were anesthetized with 2-4% isoflurane prior to placement in a stereotaxic apparatus using blunt atraumatic ear bars. Bilateral indwelling cannulas (Protech International, Roanoke, VA, C317G, D/V −6⍰mm below plate) were implanted in the vHipp (A/P −5.3 mm M/L⍰±⍰5.0 mm from Bregma D/V -6.0 mm from brain surface) and fixed in place with dental cement and four anchor screws. Rats received the analgesic ketoprofen (5 mg/kg, *s*.*c*.) and allowed to recover, individually housed, for a minimum of one week before experimentation. Injectors extending 1⍰mm past the end of the guide cannula were utilized for microinjections. Guide cannulas were kept patent with dummy cannulas. Rats were injected with either siRNA targeting RDX (0.2 ug/uL; 0.75uL) or a non-targeting, scrambled siRNA as a control at a rate of approximately 0.5 μL/min. This concentration was selected based on published data ^33^. The four RDX targeting sequences in the siRNA SMARTpool are as follows: GAAUCAGUUAUAACGUUUA; CCAAUAAAUGUAAGAGUAA; CCUUAUUGCUAAAAGAAUC; CUCUAAUUUUGGAUAAUAU. Accell siRNA (Dharmacon, Lafayette, CO, USA) was chosen specifically as it was designed to incorporate into cells that are difficult to transfect, such as neurons, without the use of a transfection agent and results in peak knockdown within 3-4 days ^34,35^.

### 2.3 Inescapable foot shock stress

Rats were randomly assigned to control (no shock) or to the shock groups that received two consecutive days of inescapable foot shock stress (IS) as previously described ^10,32^. The two-day IS paradigm consisted of placing the rats in a 30.5 × 25.4 × 30.5 cm conditioning chamber with a stainless-steel grid shock floor (Coulbourn Instruments, Whitehall, PA, USA). One session of IS consists of 60 × 15s, 0.8 mA foot shocks with an average inter-trial interval (ITI) of 30 seconds with a 25% deviation (±7.5 seconds) and lasted approximately 40 minutes. Control rats were handled daily but not exposed to conditioning chambers. Electrophysiology and behavioral experiments were conducted 24 hours after the last day of IS as previously described ^10,32^.

### 2.4 In vivo extracellular dopamine recordings

Rats were anesthetized with 8% chloral hydrate (400 mg/kg, *i*.*p*.) and placed in a stereotaxic apparatus (Kopf Instruments, Tujunga, CA, USA). Extracellular glass microelectrodes (impedance ∼6-10 MΩ) were lowered into the VTA (from bregma: AP -5.3 mm, ML ±0.6 mm, and DV -6.5 to -9.0 mm) using a hydraulic micro-positioner (Model 640, Kopf). Spontaneously active dopamine neurons were identified using open filter settings (low-frequency cutoff: 30 Hz; high-frequency cutoff: 30 kHz) according to previously established electrophysiological criteria ^36,37^. Three parameters of dopamine activity were measured and analyzed: 1.) the number of dopamine neurons firing spontaneously per track (population activity) 2.) firing rate, and 3.) proportion of action potentials occurring in bursts (defined as the incidence of spikes with <80 ms between them; termination of the burst is defined by >180 ms between spikes). Analysis of dopamine neuron activity was performed using LabChart software (ADInstruments, Sydney, Australia). Immediately following, rats were rapidly decapitated, and brains were extracted. A subset of brains was used to verify electrode and canula placement, while the remaining brains were used for molecular analysis.

### 2.5 Prepulse inhibition of startle (PPI)

Rats were placed into a sound-attenuated chamber (SD Instruments, San Diego, CA, USA) and allowed to acclimate to 65dB background noise for 5 minutes. Rats were then exposed to 10 startle-only trials [40ms, 120dB, 15s average inter-trial intervals (ITI)]. Next, rats were exposed to 24 trials where a pre-pulse (20ms at 69dB, 73dB and 81dB) is presented 100ms before the startle pulse. Each prepulse + startle pulse trial was presented 6 times in a pseudo-random order (15s average ITI). The startle response was measured from 10-80ms after the onset of the startle pulse and recorded and analyzed using SR-LAB Analysis Software (SD Instruments). PPI was calculated for each prepulse intensity and averaged across the three intensities.

### 2.6 Immunoprecipitation

Immediately following completion of electrophysiology or behavior, rats were euthanized by rapid decapitation, and the hippocampus was dissected out on ice and separated into dorsal and ventral portions. Samples were homogenized with lysis buffer and centrifuged at 14,000g for 2 minutes. Supernatants were collected and stored at -80 until α5 subunit and its binding partners were immunoprecipitated using SureBeads protein G magnetic beads according to the manufacturer’s protocol (Biorad, Hercules, CA, USA). Western blots (*detailed in section 2.8*) were used to quantify α5, RDX, and GEPH levels in both the immunoprecipitated samples and hippocampal homogenates.

### 2.7 Chemical Cross-linking Assay

A chemical cross-linking assay was performed in a subset of rats as previously described ^38,39^. Briefly, rats were rapidly decapitated, and brains extracted. The whole hippocampus was dissected out over ice, separated into dorsal and ventral portions, and minced into small pieces using a razor blade. The vHipp from one hemisphere was incubated in Dulbecco’s phosphate buffered saline (PBS) with calcium chloride and magnesium chloride (Sigma) containing bis(sulfosuccinimidyl)suberate (BS3) cross-linker (2mM, ThermoFisher, Waltham, MA, USA) for 2 hours at 4C on a shaker. The other hemisphere was incubated in Dulbecco’s PBS as a control. All samples were quenched by adding 100mM glycine and rotating another 10 minutes at 4C. Samples were then centrifuged (20,000g for 2 minutes at 4C) and supernatants were discarded. A lysis buffer containing 0.1% Triton-X-100 and peptidase inhibitors was added and tissues were homogenized (PowerGen 125, Fisher Scientific) and centrifuged for 2 minutes (20,000g at 4C). The supernatants were collected and stored at -80 until analyses by western blot.

### 2.8 Western Blot

Proteins in the lysates were separated in an SDS-PAGE gel followed by blotting onto a 0.2µm nitrocellulose membrane. Membranes were incubated with an antibody against α5GABA_A_Rs (1:1000), Radixin (1:1000), Gephyrin (1:3000), or GAPDH (1:1000) in 2.5% BSA in TBST, overnight at 4C. They were then washed with Tris-buffered saline with 0.1% Tween 20 (TBST) prior to incubation with a horseradish peroxidase-conjugated secondary antibody (goat anti-rabbit; 1: 10,000 or horse anti-mouse; 1:5000) for 1 hour at room temperature. Membranes were washed with TBST (3x for 10 min each) and incubated with a Pierce enhanced chemiluminescence kit (Thermofisher) followed by exposure to X-ray film for detection. Blots were stripped using a commercially available stripping buffer, washed, blocked, and re-probed no more than once. Densitometry analyses of immunoreactive bands were performed using the NIH Image J software from the scanned films. Densitometric arbitrary units were normalized to GAPDH, except in immunoprecipitation studies in which radixin and gephyrin measures were normalized to α5GABA_A_R levels.

### 2.9 Histology

To verify electrode and cannula placement, brains were fixed for at least 24 hours (4% phosphate buffered formaldehyde), and cryoprotected (10% w/v sucrose in phosphate-buffered saline) until saturated. Brains were coronally sectioned (25 µm) using a cryostat (Leica, Buffalo Grove, IL, USA). Sections containing electrode or cannula tracks were mounted onto gelatin-coated slides, stained with neutral red (0.1%) and thionin acetate (0.01%), and cover-slipped with DPX Mountant for histochemical confirmation within the VTA (electrode) or vHipp (cannula) (**Figure 2A**).

**Figure 2.**
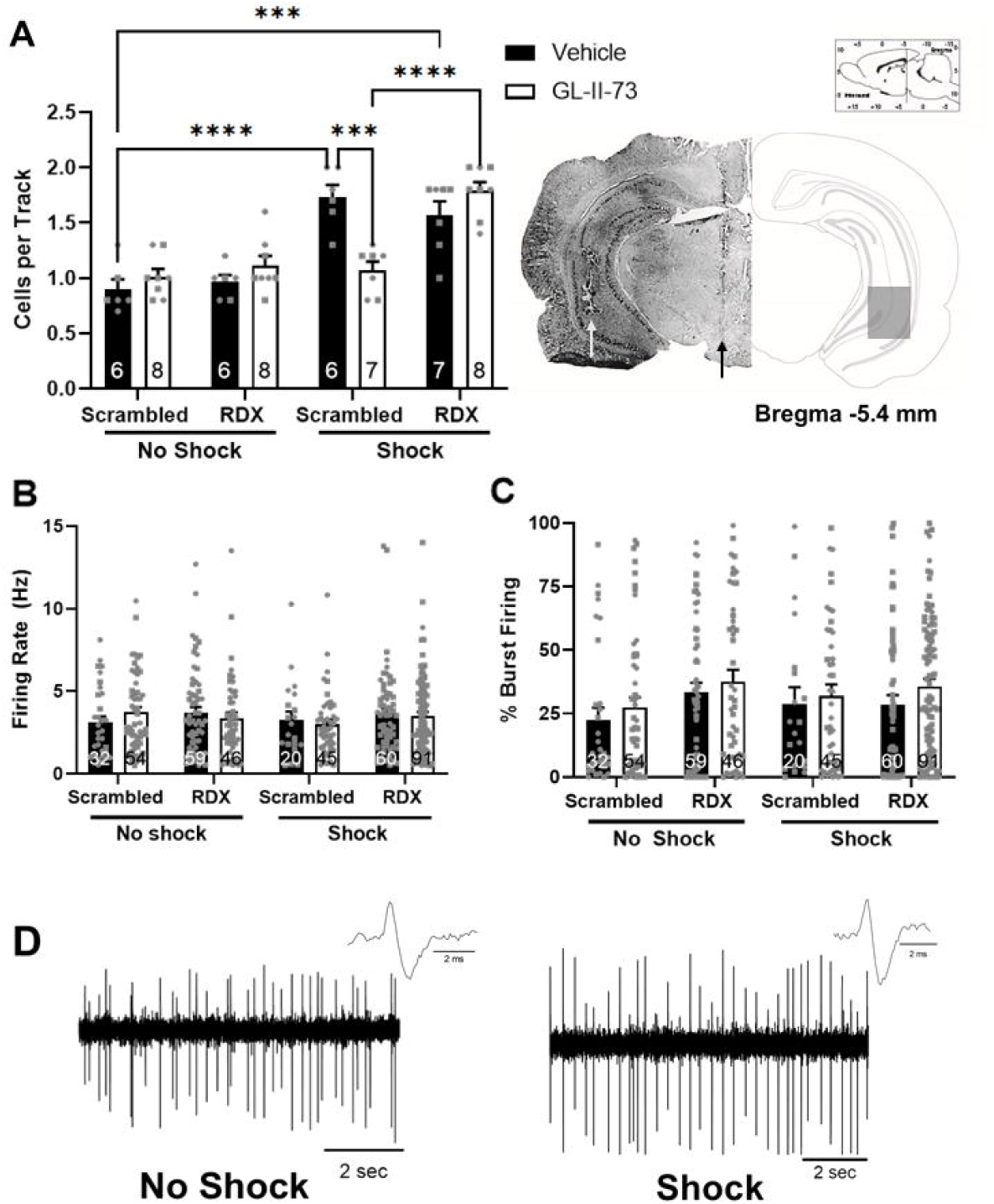
GL-II-73 was unable to restore dopamine system function when radixin is knocked down. *In vivo extracellular* electrophysiology was used to measure dopamine cell activity in the ventral tegmental area. **A**) *Left*, Inescapable shock exposure significantly increased the number of spontaneously active cells/track (population activity), which was reversed by intra-ventral hippocampus injection of GL-II-73 (100ng/μL; 0.75 μL), but not when radixin was knocked down. *Right*, Representative brain slice with electrode placement in the ventral tegmental area (VTA) (black arrow) and cannula placement for drug administration in the ventral hippocampus (vHipp, white arrow), with corresponding schematics of the brain section (−5.40 mm posterior to bregma) with box indicating the area in which tracks were found. Neither (***B***) firing rate nor (***C***) burst firing was affected by siRNA, shock, or drug treatment. (***D***) representative traces from control (left) and shocked (right) rats. n=6-8/group, males and females represented as circles and squares, respectively \*\*\**P* = 0.0001, \*\*\*\**P* < 0.0001, RDX = Radixin

### 2.10 Materials

The proprietary compound, GL-II-73, was synthesized by the University of Wisconsin-Milwaukee and supplied by the Centre for Addiction and Mental Health, Campbell Family Mental Health Research Institute (Toronto, ON, CA),. Chloral hydrate (C8383), propylene glycol (P4347), and Tween80 (P1754) were obtained from Sigma-Aldrich (St. Louis, MO, USA). Antibodies were from R&D Systems, (Minneapolis, MN USA,) #PPS027 (α5) or Abcam (Cambridge, UK) ab5249 (radixin) ab181382 (gephryin) #9484 (GAPDH) or Cell signaling (Davers, MA, USA) #7074 (anti-rabbit-HRP) #7076 (anti-mouse-HRP). Accell siRNA were purchased from Dharmacon.

### 2.11 Statistical analysis

Data are represented as mean⍰±⍰SEM and n values representing either the number of rats or neurons as indicated. In all experiments, data were analyzed by three-way ANOVA (electrophysiology and PPI; factors: stress x drug x siRNA), two-way ANOVA (α5 surface expression; factors: siRNA x crosslinker) or t-test (Western Blot) and plotted using Prism software (GraphPad Software Inc.; San Diego, CA, USA). When significant main effects or interactions were detected the Holm–Sidak post-hoc test was used. All tests were two-tailed, and significance was determined at p⍰<⍰0.05. While both sexes were represented, we were not powered to detect sex differences and therefore did not explicitly test for this. Raw electrophysiology data were analyzed using LabChart version 8 (ADInstruments, Colorado Springs, CO, USA) and PPI data were analyzed using SR Labs Analysis software (SD Instruments).

## 3. Results

### 3.1 The therapeutic effects of intra-vHipp administration of GL-II-73 on dopamine system function are blocked by radixin knockdown

To evaluate dopamine system function, we measured dopamine neuron activity in the VTA using *in vivo* extracellular electrophysiology. Consistent with previous findings ^10,32^, inescapable footshock stress elicited a significant increase in population activity (n= 6 rats; 1.733⍰±⍰0.109 cells per track; three-way ANOVA; *F*_*Shock(1,48*)_ = 73.860; *p* <⍰0.0001; *F*_*siRNA(1,48*)_= 8.135; *p* =⍰0.006; Holm–Sidak; *t*⍰=⍰ 6.153, *p*⍰<⍰0.0001; **Figure 2A**) when compared to non-shocked vehicle rats (n=6 rats; 0.900⍰±⍰0.089 cells per track). This shock-induced increase in dopamine neuron activity was completely reversed by intra-hippocampal administration of GL-II-73 (n= 7 rats; 1.071⍰±⍰0.078 cells per track; Holm–Sidak; *t* = 5.072, *p*⍰=⍰0.0001) and had no effect in non-shocked rats who received intra-hippocampal GL-II-73 (n= 8 rats; 1.013⍰±⍰0.069 cells per track). Further, knocking down radixin in non-shocked rats had no effect on population activity in vehicle (n= 6 rats; 0.967⍰±⍰0.061 cells per track) and GL-II-73 treated rats (n= 8 rats; 1.113⍰±⍰0.088 cells per track). Again, consistent with observations in rats who received the scrambled siRNA, shock produced a significant increase in dopamine neuron activity, (n= 7 rats; 1.571⍰±⍰0.121 cells per track; Holm–Sidak; *t* = 1.780, *p*⍰=⍰0.669) in radixin knockdown rats. Interestingly, increasing synaptic α5GABA_A_R localization by knocking down radixin blocked the ability of intra-hippocampal GL-II-73 to restore dopamine system function in shocked rats (n=8 rats; 1.788⍰±⍰0.081 cells per track). As expected, no significant differences were observed in the average firing rate (**Figure 2B**; *F*_Shock (*1,404*)_ = 0.953; *p* = 0.330; *F* _siRNA (*1,404)*_ =0.304; *p* = ⍰0.582 *F*_drug (*1,404)*_ = 0.033; *p* = ⍰0.856) or burst firing (**Figure 2C**; *F*_Shock (*1, 404)*_ =3.427; *p* =⍰0.065; *F* _siRNA (*1, 404)*_= 0.044; *p* =⍰0.833; *F* _drug (*1, 404)*_= 2.200; *p* = ⍰0.139). Representative traces from control (left) and shock (right) animals are shown in **Figure 2D**.

### 3.2 Increased synaptic α5GABAAR expression does not prevent the effects of GL-II-73 in prepulse inhibition of startle

To evaluate sensorimotor gating, we measured prepulse inhibition of the acoustic startle response **(Figure 3)**. Previous studies measuring PPI in rats exposed to IS reported a significant decrease in the %PPI following inescapable footshock stress ^10,32^. Here, we observed a significant main effect of shock (n= 9-11 rats per group; three-way ANOVA; *F*_shock (1, 71)_ = 14.310; *p* = 0.0003) and of intra-vHipp administration of GL-II-73 (*F*_drug (1, 71)_ = 8.765; *p*=0.004); however, *post hoc* tests revealed no significant differences between groups.

**Figure 3.**
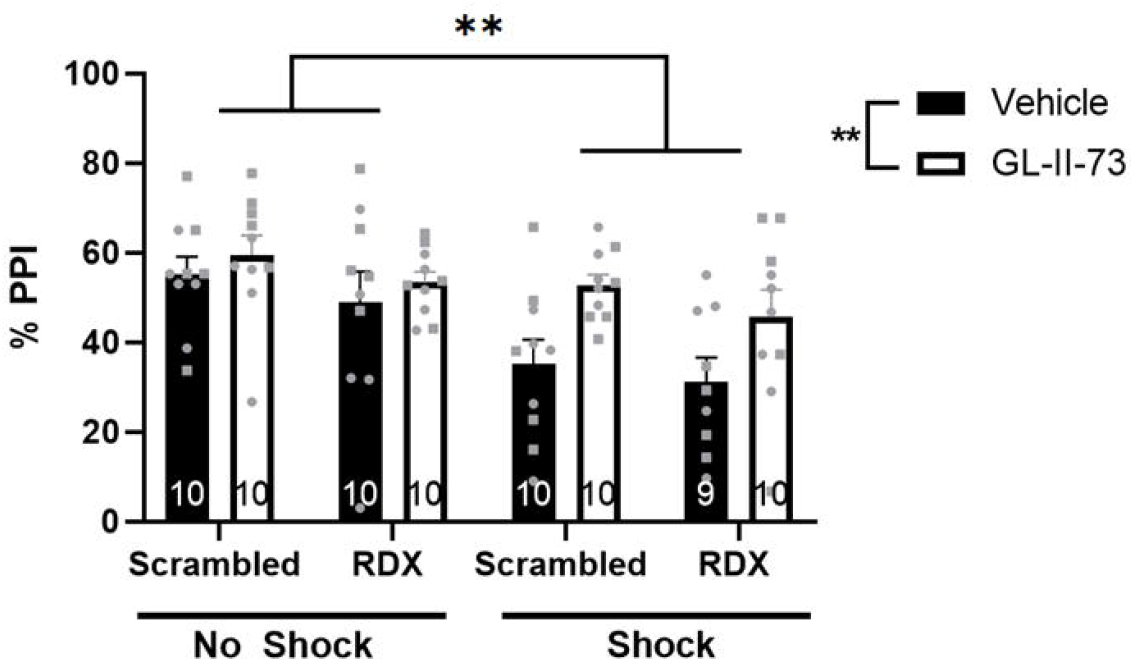
Radixin knockdown does not alter prepulse inhibition. Two days of inescapable shock had a significant main effect on PPI as did treatment with GL-II-73 (p = 0.0042), however, post hoc analysis revealed no relevant group differences. n=9-11/group, males and females represented as circles and squares, respectively. \*\**P* < 0.005, RDX = radixin.

### 3.3 Radixin knockdown increased markers of synaptic α5GABAAR but did not alter α5GABAAR expression within the vHipp

To validate the successful knockdown of radixin, we measured radixin associated with α5GABA_A_R using coimmunoprecipitation in control rats. We observed a significant difference between radixin knockdown and control groups (**Figure 4A**, t test; *t* = 2.629, *p* = 0.017). Additionally, co-immunoprecipitation of α5GABA_A_R and gephyrin revealed a significant increase in gephyrin levels in rats that had radixin knocked down compared to controls (**Figure 4B**, t test; *t*=3.069, *p*=0.008), suggesting an increase in synaptic α5GABA_A_Rs. Finally, to ensure any electrophysiology and behavioral results were not due to degradation or internalization of α5GABA_A_Rs when radixin is knocked down, we also measured surface and total α5GABA_A_R using a chemical crosslinking assay. While there was an expected significant difference between crosslinked samples and homogenate (**Figure 3C**, two-way ANOVA, *F*_crosslinker (1, 16)_ = 11.300; *p* = 0.004) there were no significant differences due to radixin knockdown (*F*_siRNA (1, 16)_ = 0.173; *p* = 0.683).

**Figure 4:**
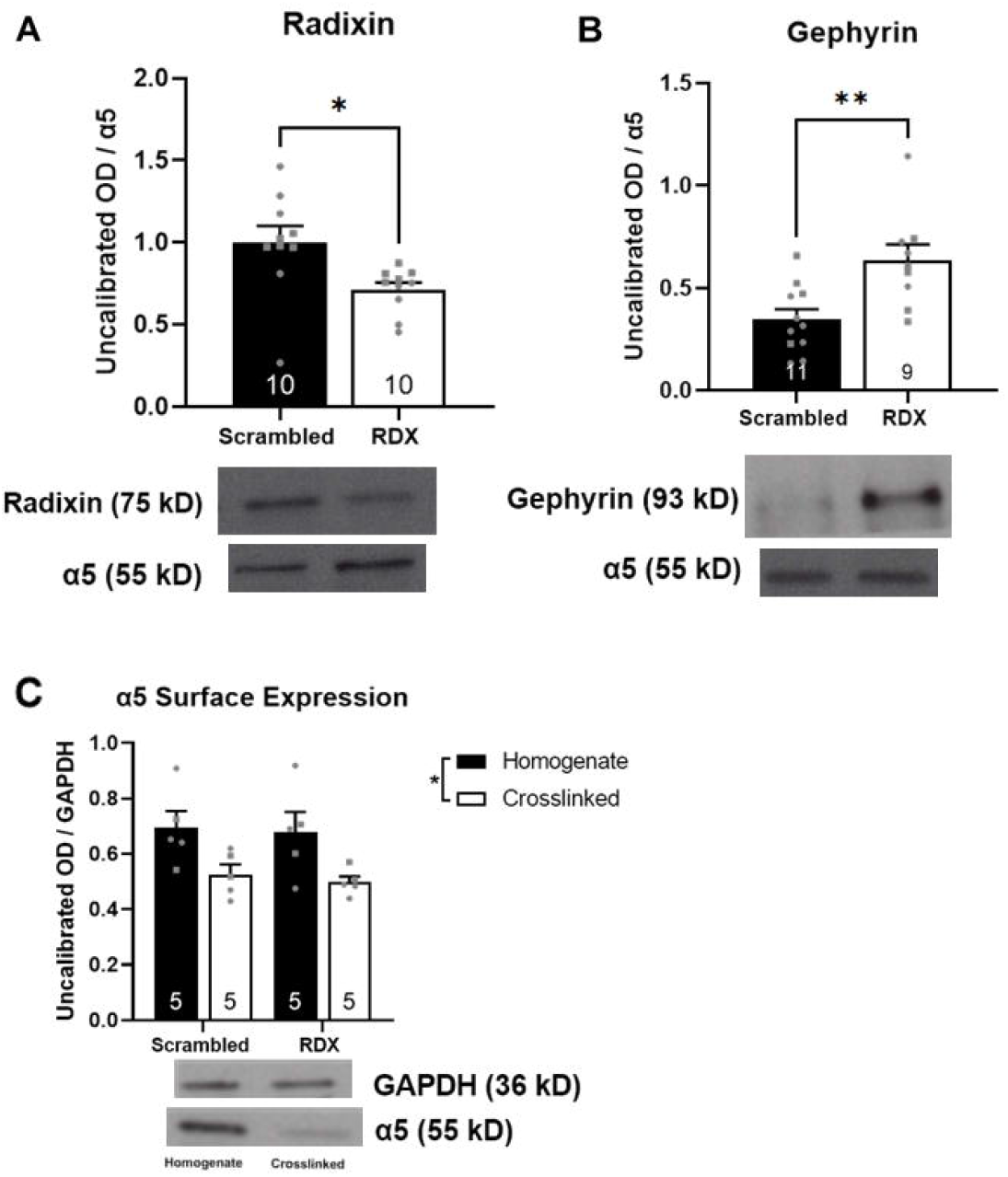
Radixin knockdown increases synaptic α5, without changing surface or total α5 expression. **A)** Co-immunoprecipitation of α5 and radixin revealed a significant decrease in α5-associated radixin in rats that received radixin-targeted siRNA compared to those that received scrambled siRNA. Representative image of bands below. n=10/group. **B**) Conversely, co-immunoprecipitation of α5 and gephyrin revealed a significant increase in gephyrin levels in the radixin knockdown group. Representative image of bands below. n=9-11/group **C)** Treatment with the crosslinking agent caused a significant decrease in optical density of α5 immunoreactive bands, but no differences in total α5 (homogenate) or surface (crosslinked) were observed between rats that received scrambled siRNA or radixin-targeted siRNA. Representative image of bands below graphs. n=5/group, males and females represented as circles and squares, respectively. *p<0.05, **p<0.01 n = 10, RDX = Radixin, OD = optical density.

## Discussion

Psychosis is debilitating symptom that accompanies many neurological disorders, including PTSD ^31,40^. The dopamine hypothesis states that aberrant dopamine neuron activity underlies psychosis symptoms, yet currently available antipsychotics that target dopamine D2 receptors are not always effective and often result in intolerable side effects (i.e. dyskinesias and metabolic disorders) ^41^. This has led some to suggest that indirectly modulating dopamine neuron activity through manipulating activity in upstream brain regions may be an effective treatment strategy that produces fewer adverse effects. The hippocampus is a brain region that can modulate dopamine neuron activity through a multisynaptic pathway starting in the nucleus accumbens ^12,13^. Using a multitude of techniques, we and others have demonstrated that attenuating vHipp activity can restore dopamine neuron population activity and related behaviors in animal models used to study psychosis ^12,16,18,19,42^. An effective and translational approach to inhibiting vHipp activity is by using α5-PAMs. Indeed, we and others have previously shown that targeting α5GABA_A_Rs can improve physiological and behavioral alterations associated with psychosis ^9–11,20^, suggesting that α5-PAMs may possess antipsychotic efficacy. Interestingly, the efficacy of PAMs appears to be specific to those selective for the α5-subunit, as targeting vHipp α1GABA_A_Rs does not appear to modulate VTA dopamine neuron activity ^11,20^. A major delineation between these receptor types is their cellular location, with α5GABA_A_Rs existing both in the synapse and extrasynaptic space whereas α1GABA_A_Rs are limited to the synapse. Here, we examined if the observed differences in antipsychotic-like efficacy were due to receptor location (extrasynaptic vs synaptic). Based on our previous findings with targeting synaptic α1GABA_A_Rs, we posited that GL-II-73 would no longer be able to modulate dopamine neuron activity when α5GABA_A_Rs are shifted into the synapse. These studies have important implications, as previous studies have determined that high periods of hippocampal activity, often observed in psychosis ^12,13,15^, can promote movement of α5GABA_A_Rs into the synapse. Aberrant dopamine neuron activity is central to the pathology of psychosis and is observed in both patients ^43–45^ and rodent models ^46^. To examine dopamine system function, we use *in vivo* electrophysiology to measure the number of spontaneously active dopamine neurons in the VTA, referred to as population activity ^14^. We consistently find that animal models used to study psychosis have elevated dopamine neuron activity ^17,19,47^. Here, we report that IS exposure, a common rodent model to study PTSD, induced aberrant dopamine neuron population activity, a finding consistent with previous literature ^10,32^. This was reversed by GL-II-73. However, in conditions of knocking down radixin, which caused a shift of α5GABA_A_Rs to the synapse, this effect of GL-II-73 was lost. These results suggest that the ability of GL-II-73 to modulate VTA dopamine neuron activity is dependent on the extrasynaptic localization of α5GABA_A_Rs.

Patients with PTSD and patients with psychosis both display deficits in sensorimotor gating ^48–50^, a behavioral dimension that is readily assessed in rodents using PPI ^51^. Indeed, rodent models used to study both PTSD and psychosis display deficits in PPI ^32,52^, which can be reversed by GL-II-73 ^10^. In the current study we demonstrated that IS decreases PPI and that intervention with GL-II-73 attenuates this, regardless of the radixin knockdown (i.e., regardless of the localization of α5GABA_A_Rs). While PPI is a dopamine-dependent behavior, it is mediated by other circuits and not exclusively controlled by dopamine^29^. This may suggest that the reliance of GL-II-73 on extrasynaptic receptors is limited to modulation of dopamine neuron activity, and may not apply to the antidepressant-like effects ^6^ or the pro-cognitive effects ^7^. Nonselective benzodiazepines, or derivatives that primarily act on α1GABA_A_Rs are ineffective as antipsychotics ^53,54^. This is in line with our previous studies demonstrating that selectively targeting α1GABA_A_Rs or nonselectively targeting GABA_A_Rs in the vHipp does not affect dopamine neuron activity ^11,55^. Taken with the findings presented in the current study, it appears that this is due to targeting of synaptic GABA_A_Rs. However, an open question remains as to why synaptic GABA_A_Rs do not modulate VTA dopamine neuron activity in the way that extrasynaptic ones can. One possibility is that the type of inhibition produced by extrasynaptic receptors (tonic) is more effective at maintaining a decrease in hippocampal activity than synaptic receptors (phasic). It is possible that the fast, transient nature of phasic inhibition is insufficient to produce changes in downstream dopamine activity, whereas the relatively slower and more persistent effects of α5-mediated tonic inhibition has a more robust effect ^56,57^. Although this explanation only partially accounts for earlier studies which demonstrate that dampening excitatory transmission in the vHipp using tetrodotoxin can also restore healthy dopamine system function in animal models used to study psychosis ^16,42,58^.

The loss of efficacy when moved into the synapse may also be explained by a change in receptor functionality. While it has been shown that synaptic α5GABA_A_Rs can successfully contribute to IPSCs ^24,27,28^, differences in structure caused by loss or gain of protein-protein interactions may prevent GL-II-73 from modulating α5GABA_A_R when they move to the synapse. For example, it is known that α5GABA_A_Rs interact with auxiliary subunits, such as Shisa7, which can modify receptor kinetics ^59^ and trafficking ^60^ and appear to be critical for tonic currents ^61^. Alterations in protein interactions may change receptor function enough to negate the effects of GL-II-73 in the situation of α5GABA_A_Rs being localized to the synapse.

We confirmed that decoupling α5GABA_A_Rs from radixin did not reduce membrane expression of α5GABA_A_Rs, suggesting that the absence of an effect of GL-II-73 in radixin knockdown rats is not due to a reduction in receptor availability. However, a limitation of this study is that we did not measure actual levels of synaptic and extrasynaptic receptors. Rather, we measured markers of α5GABA_A_Rs localization through association with radixin and gephyrin. Thus, while our coimmunoprecipitation studies suggest that knockdown of radixin increases synaptic α5GABA_A_Rs, we acknowledge the caveat associated with measuring proxies for localization. Future studies should more rigorously examine the dynamics of α5GABA_A_R relocalization and pinpoint the biological processes resulting in the dramatic loss of efficacy we observed here as this may have important clinical implications, especially as interest in α5-PAMs from the pharmaceutical industry increases. Indeed, α5GABA_A_R localization appears to be dynamically modulated by hippocampal activity levels^24^. It is possible that certain conditions where hippocampal activity is dramatically altered (e.g., epilepsy-induced psychosis) the proportion of synaptic α5GABA_A_Rs could increase, diminishing the ability of GL-II-73 and perhaps other α5-PAMs as well. Although the results obtained here suggest that this is limited to modulation of dopamine neuron activity, as PPI was unaffected by radixin knockdown. This study highlights the importance of testing novel therapeutics in multiple disease states and/or models, a concept of particular importance for GL-II-73, which has shown promising therapeutic potential for a variety of psychiatric conditions ^6,7,10,11^.

## Author Contributions

A.M.M. made contributions to the design of the work, acquisition, analysis and interpretation of the data, as well as drafting and editing the manuscript. T.D.P. and E.L.S. assisted in interpretation of the data and revising and editing the manuscript content. M.Y.M. and D.S. synthesized GL-II-73 under the supervision of J.C. A.N.A. contributed to acquisition of the data. D.J.L. contributed to the concept and design of the study, analysis and interpretation of the data, revising and editing manuscript content, as well as providing final approval of the document.

